# Somatic mutation of Afadin leads to anchorage independent survival and metastatic growth of breast cancer through αE-catenin dependent destabilization of the adherens junction

**DOI:** 10.1101/2023.07.04.547654

**Authors:** Max A.K. Ratze, Lotte N.F.L. Enserink, Noboru Ishiyama, Christina H.J. Veltman, Isaac J. Nijman, Rene Bernards, Paul J. van Diest, Matthias Christgen, Patrick W.B. Derksen

**Author notes:** **Correspondence to:** Patrick W.B. Derksen Department of Pathology, H04.312 Heidelberglaan 100, 3584CX, Utrecht, The Netherlands.

## Abstract

Loss of E-cadherin (*CDH1*) and the adherens junction (AJ) drive development and progression of invasive lobular breast cancer (ILC). However, approximately 40% retain wild type *CDH1* alleles, indicating that modulation of other genes attenuates the AJ during ILC etiology. To identify alternative drivers, we performed targeted sequencing in *CDH1* wild type samples, based on a defined set of 100 AJ, tight junction, and desmosome genes we designated as the ‘Adhesome’. In 146 ILC samples, we identified 62 cases (43%) with wild type *CDH1* alleles in which we detected a total of 284 mutations in 36 Adhesome genes. After selection based on occurrence and potential loss of function, we identified an inactivating frameshift mutation in Afadin (*AFDN; p*.Lys630fs).

Functional studies in E-cadherin-expressing breast cancer cells showed that Afadin knockout leads to immature AJs, and a non-cohesive phenotype accompanied by actomyosin dependent anoikis resistance, which are classical ILC hallmarks. Afadin reconstitutions show that F-actin organization critically depends on the ⍰E-catenin binding CC domain. Afadin loss in intraductal xenograft mouse breast cancer models leads to ILC-type morphologies and overt lung metastases. *AFDN* truncate reconstitutions revealed that deletion of the C-terminal ⍰E-catenin binding CC domain is sufficient to drive metastatic ILC. In conclusion, we identified and functionally coupled a somatic frameshift *AFDN* mutation in breast cancer to destabilization the epithelial AJ and the development of ILC hallmarks such as actomyosin-dependent anoikis resistance and single cell invasion. As such, Afadin represents a candidate tumor suppressor for E-cadherin-positive ILC development and progression.

## INTRODUCTION

Invasive lobular carcinoma (ILC) constitutes up to 15% of all breast cancers, making it the second most common breast cancer type. ILC is characterized by non-cohesive growth patterns and diffuse dissemination to specific sites such as the gastrointestinal tract, ovaries and peritoneum (Rakha & Ellis, 2010). ILC development and progression are caused by a functional inactivation of E-cadherin (*CDH1*) (Berx *et al*, 1995; Derksen *et al*, 2011, 2006), a transmembrane molecule that represents the principal component of the adherens junction (AJ) (Meng & Takeichi, 2009). A general hallmark of ILC cells is the acquisition of anchorage independent survival, also called anoikis resistance (Christgen & Derksen, 2015). Anoikis resistance in ILC cells is primarily driven by an autocrine activation of growth factor receptors and their effectors such as PI3K/AKT/FOXO/BMF (Hornsveld *et al*, 2016; Nagle *et al*, 2018; Teo *et al*, 2018), and constitutive activation of Rock1-dependent actomyosin contraction (Schackmann *et al*, 2011; Ven *et al*, 2015). Notwithstanding this, it is well established that approximately 10% of ILC cases preserve membranous E-cadherin expression (Reed *et al*, 2016; Grabenstetter *et al*, 2020). Moreover, somatic inactivating mutations in *CDH1* do not fully account for the observed E-cadherin loss in ILC, as approximately 30-40% retain the wild-type alleles (Ciriello *et al*, 2015; Desmedt *et al*, 2016; Pareja *et al*, 2020). These findings imply that alternative drivers may exist beyond *CDH1* that confer an ILC tumor suppressor function.

The epithelial AJ is essential for cell-cell connections between two neighboring cells. Extracellular E-cadherin homotypic interactions *in cis* and *in trans* provide a mechanical signaling hub within the AJ through linkage to the actin cytoskeleton (Yamada *et al*, 2005; Nelson, 2008; Kurita *et al*, 2011). Upon cell-cell contact, the cytosolic E-cadherin tail interacts with p120-catenin (*CTNND1*), β-catenin (*CTNNB1*) and αE-catenin (*CTNNA1,* α-catenin from hereon), proteins that are essential for the stability of the AJ (Anastasiadis *et al*, 2000; Davis *et al*, 2003) and for mechanical and biochemical signal transduction towards F-actin respectively (reviewed in: (Leckband & Rooij, 2014)). Linking of the cytoskeleton occurs by indirect binding through either classical Cadherins or Nectins via their associated accessory proteins. Connections to the F-actin cytoskeleton are dependent on binding to β-catenin (*CTNNB1)* and the subsequent interaction with α-catenin, facilitating mechanically transduced cues via the F-actin cytoskeleton and its dynamics (Meng & Takeichi, 2009; Shapiro & Weis, 2009; Efimova & Svitkina, 2018). Interestingly, interaction of α-catenin with the F-actin cytoskeleton is dependent on interactions with additional proteins such as Vinculin and Eplin to build up the initial forces along the F-actin cytoskeleton and modulate the actin cytoskeleton in response to forces applied to the AJ (Weiss *et al*, 1998; Dufour *et al*, 2013; Collins *et al*, 2015). Interestingly, α- catenin has been advocated as an alternative mode for tumor suppression in malignancies that are caused by E-cadherin dysfunction, such as ILC and diffuse gastric cancer (Majewski *et al*, 2013; Blair *et al*, 2020). Functional follow-up studies in breast cancer subsequently showed that α-catenin loss indeed leads to ILC-like phenotypes in culture and in mice, and acquisition of actomyosin-dependent anoikis resistance (Hollestelle *et al*, 2010; Groot *et al*, 2018). Together, these findings indicate that loss of components that function in cell-cell contacts (other than E-cadherin) can induce destabilization of the AJ and underpin subsequent tumor etiology.

E-cadherin and Nectin both control linkage to the F-actin cytoskeleton. Unlike E-cadherin, Nectin mediates F-actin modulation primarily through Afadin (*AFDN* or *MLLT4*) (Mandai *et al*, 1997; Rikitake & Takai, 2011) (Reviewed in: (Huxham *et al*, 2021)). The longer l-Afadin splice variant (hereafter Afadin) controls these connections as a multi-domain scaffolding protein that interacts with various tight junction (TJ) components, indirect connections through α-catenin and the AJ (Sakakibara *et al*, 2020), or direct interaction with F-actin via its Coiled Coil (CC) and/or F-Actin Binding (FAB) domains (Mandai *et al*, 1997; Choi *et al*, 2016; Carminati *et al*, 2016). Moreover, binding of *AFDN* with LMO7 and ADIP allow for interaction of Nectin-based junctions with Cadherin-based AJs (Asada *et al*, 2003; Ooshio *et al*, 2004). Its favorable localization due to its function to link the F-actin cytoskeleton to the nectin-based AJ, enables it to support the connection between the F-actin cytoskeleton and the AJ (Takai *et al*, 2008; Sakakibara *et al*, 2020).

Cell-cell interactions are additionally reinforced by the formation of TJs, multiprotein junctional complexes that are essential in the formation of a functional barrier in epithelium and endothelium and correct apical polarization of cells (Zihni *et al*, 2016). Like the AJ, TJs facilitate cell-cell interactions through a complex arrangement of transmembrane proteins that interact cytoplasmic adaptors such as zona occludens proteins (Morita *et al*, 1999). Zona occludens 1 (ZO1) links the TJ to the AJ through complex formation with α-catenin and recruitment of JAMA to the Nectin-based junctions (Fukuhara *et al*, 2002; Maiers *et al*, 2013). As such, proteins within the TJ may be critical to the tumor suppressive function of the AJ. Indeed, inactivation of the TJ polarity protein PAR3 induces destabilization of AJs, leading to disrupted actin dynamics and invasion of breast cancer cell (McCaffrey *et al*, 2012; Xue *et al*, 2013).

Here, we have performed targeted sequencing on ILC samples to reveal alternative drivers of ILC. In a cohort of clinical samples that had retained wild type *CDH1* alleles, we identify Afadin as a candidate tumor suppressor for the development and progression of ILC. Functionally, loss of Afadin induces an E-cadherin positive ILC phenotype, and tumor progression through the acquisition of anoikis resistance, invasion, and metastatic capacity.

## RESULTS

### Identification of alternative driver mutations in *CDH1* wild-type ILC

To identify alternative drivers of ILC, we performed a focused gene set exome sequencing analysis on 86 genes that play an essential role in epithelial cell-cell adhesion. This gene set (from hereon: the Adhesome) contains genes coding for AJ, TJ and desmosome (DS) proteins (Table 1). In total, 154 ILC samples from the RATHER cohort (Michaut *et al*, 2016) were sequenced using Adhesome-targeted Next-Generation Sequencing, of which 146 samples passed the sequencing quality checks (Fig. 1A). Deleterious *CDH1* mutations were found in 84 samples (57.5% of total), which is in line with previously published numbers for *CDH1* status in ILC (Berx *et al*, 1995; Pareja *et al*, 2020). To enable identification of alternative ILC drivers, *CDH1* mutant samples were excluded. Also, mutations that occurred more than three times were considered nucleotide polymorphisms and were omitted from further analysis. In the remaining 62 samples, 284 non-synonymous mutations were identified in a total of 78 different genes (Fig. 1B). We next focused on mutations with a potential major impact on loss of protein function (start lost/stop gained, frameshift and missense mutations), which led to the identification of Afadin (*AFDN*), which we prioritized based on its published established roles in cell-cell adhesion. In this ILC sample, *AFDN* contained a heterozygous p.Lys630Stop frameshift mutation that potentially leads to truncation and loss of Afadin protein expression (Fig. 1C). Next, we examined the corresponding primary ILC sample (R-L1047-C), which featured small to medium-sized breast cancer cells arranged in one-to two-cell thick trabeculae and single cell files (Fig. 1D). Foci with dissociative growth were also evident. Mammary ducts were encircled by tumor cells in a targetoid fashion and tumor cells accumulated at the border from connective to adipose tissue. Tumor cells showed scanty cytosol, delicate nuclear chromatin and focal biconcave nuclear compressions. Mitotic activity was low. The final diagnosis was ILC grade 2. Importantly, using immunofluorescence, we observed that Afadin expression was lost in this sample, accompanied by a relatively weak and non-uniform membranous expression of E-cadherin (Fig. 1E).

**Figure 1.**
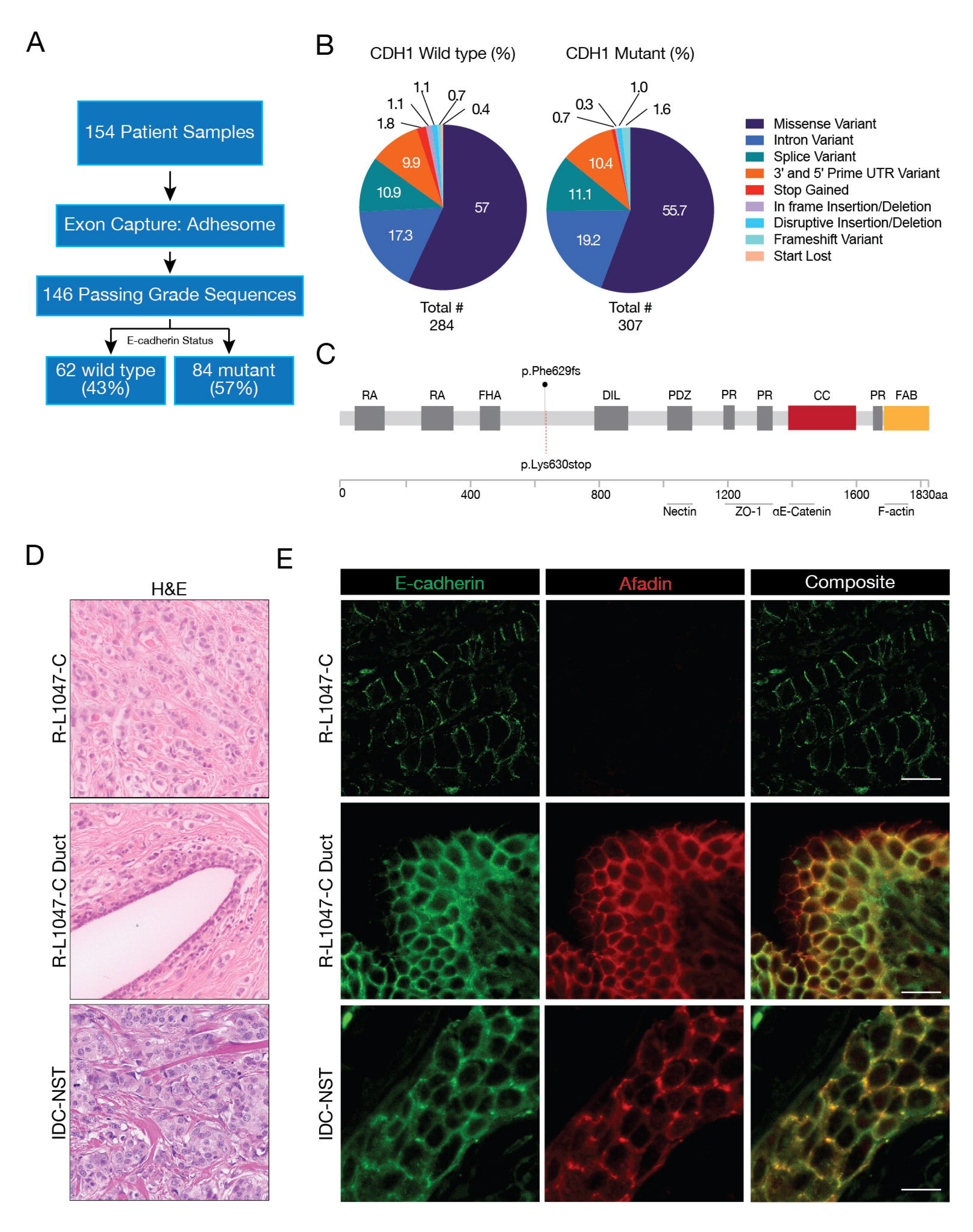
Identification of Adhesome mutations in *CDH1* wild-type patients. **A.** Dataflow for the Adhesome analysis in ILC. Indicated are the percentages *CDH1* wild type *versus* mutant ILC cases. **B.** Distribution of Adhesome mutation types. Pie charts depicting the mutation types found in *CDH1* wild-type and mutant ILC. Numbers represent % of total. **C.** Schematic depiction of the Afadin protein. Indicated are the functional domains, the identified frameshift mutation, and the predicted truncating stop codon in patient R-L1047-C. Domains and regions: RA, Ras associated; FHA, forkhead associated; DIL, Dilution; PR, proline rich; CC, coiled coil; FAB, F-actin binding. The binding sites for the effector proteins are indicated. Partly adapted from (Mandai *et al*, 2013) **D.** Histology from sample R-L1047-C (top), a healthy mammary duct with ILC infiltrate from sample R-L1047-C (middle), accompanied by a representative Invasive Ductal Carcinoma-Non Specific Type (IDC-NST) sample (bottom) **E.** Afadin protein expression is lost in ILC sample R-L1047-C containing the identified *p.*Phe629fs mutation. Shown are immunofluorescence images for E-cadherin (left) and Afadin (middle) expression. Shown are ILC sample R-L1047-C (top), a healthy mammary duct with ILC infiltrate from sample R-L1047-C (middle), accompanied by a representative IDC-NST sample (bottom). Right panels show the merged composite. Size bar = 100 µM.

**Table 1.** The Adhesome full gene list.

| Name | symbol | complex | Name | symbol | complex |
| --- | --- | --- | --- | --- | --- |
| 1 α-catenin | CTNNA1 | AJ | 44 γ-catenin | JUP | DS |
| 2 Afadin | MLLT4 | TJ | 45 Hakai | CBLL1 | AJ |
| 3 Ajuba | JUB | AJ | 46 JAM-2 | JAM2 | TJ |
| 4 aPKC | PRKCA | TJ | 47 JAM-3 | JAM3 | TJ |
| 5 Axin | AXIN1 | AJ | 48 JAM-A | F11R | TJ |
| 6 β-catenin | CTNNB1 | AJ | 49 Kaiso | ZBTB33 | AJ/OTHER |
| 7 CD151 | CD151 | OTHER | 50 KIF1C | KIF1C | OTHER |
| 8 Cingulin | CGN | TJ | 51 KIF4A | KIF4A | OTHER |
| 9 Claudin-1 | CLDN1 | TJ | 52 KIF14 | KIF14 | OTHER |
| 10 Claudin-10 | CLDN10 | TJ | 53 KIF23 | KIF23 | AJ/OTHER |
| 11 Claudin-11 | CLDN11 | TJ | 54 KIFC3 | KIFC3 | AJ/OTHER |
| 12 Claudin-12 | CLDN12 | TJ | 55 NEZHA | CAMSAP3 | AJ/OTHER |
| 13 Claudin-14 | CLDN14 | TJ | 56 MRIP | MPRIP | OTHER |
| 14 Claudin-15 | CLDN15 | TJ | 57 MUPP-1 | MPDZ | TJ |
| 15 Claudin-16 | CLDN16 | TJ | 58 Nectin-1 | CADM3 | OTHER |
| 16 Claudin-17 | CLDN17 | TJ | 59 Nectin-2 | CADM1 | OTHER |
| 17 Claudin-18 | CLDN18 | TJ | 60 Nectin-3 | CADM2 | OTHER |
| 18 Claudin-19 | CLDN19 | TJ | 61 Nectin-4 | CADM4 | OTHER |
| 19 Claudin-2 | CLDN2 | TJ | 62 occludin | OCLN | TJ |
| 20 Claudin-20 | CLDN20 | TJ | 63 P-cadherin | CDH3 | AJ |
| 21 Claudin-22 | CLDN22 | TJ | 64 p120 | CTNND1 | AJ |
| 22 Claudin-23 | CLDN23 | TJ | 65 PAR3 | PARD3 | TJ |
| 23 Claudin-24 | CLDN24 | TJ | 66 PAR6 | PARD6A | TJ |
| 24 Claudin-25 | CLDN25 | TJ | 67 PATJ | INADL | TJ |
| 25 Claudin-3 | CLDN3 | TJ | 68 Perp | PERP | TJ |
| 26 Claudin-4 | CLDN4 | TJ | 69 PI3K/p110 | PIK3CA | AJ/OTHER |
| 27 Claudin-5 | CLDN5 | TJ | 70 Plakophilin-1 | PKP1 | DS |
| 28 Claudin-6 | CLDN6 | TJ | 71 Plakophilin-2 | PKP2 | DS |
| 29 Claudin-7 | CLDN7 | TJ | 72 Plakophilin-3 | PKP3 | DS |
| 30 Claudin-8 | CLDN8 | TJ | 73 Plakophilin-4 | PKP4 | DS |
| 31 Claudin-9 | CLDN9 | TJ | 74 PLEKHA7 | PLEKHA7 | AJ |
| 32 Desmocollin-1 | DSC1 | DS | 75 Scribble | SCRIB | TJ |
| 33 Desmocollin-2 | DSC2 | DS | 76 Talin | TLN1 | OTHER |
| 34 Desmocollin-3 | DSC3 | DS | 77 TAZ | WWTR1 | OTHER |
| 35 Desmoglein-1 | DSG1 | DS | 78 Tricellulin | MARVELD2 | TJ |
| 36 Desmoglein-2 | DSG2 | DS | 79 VASP | VASP | OTHER |
| 37 Desmoglein-3 | DSG3 | DS | 80 vinculin | VCL | AJ |
| 38 Desmoglein-4 | DSG4 | DS | 81 N-WASP | WASL | OTHER |
| 39 Desmoplakin | DSP | DS | 82 YAP | YAP1 | OTHER |
| 40 E-cadherin | CDH1 | AJ | 83 ZO-1 | TJP1 | TJ |
| 41 ENA | ENAH | AJ/OTHER | 84 ZO-2 | TJP2 | TJ |
| 42 Eplin | LIMA1 | AJ | 85 ZO-3 | TJP3 | TJ |
| 43 Fer | FER | AJ/OTHER | 86 Zyxin | ZYX | AJ/OTHER |

AJ=Adherens Junction; TJ=Tight Junction; DS=Desmosome

### Somatic *AFDN* mutations are associated with *CDH1* wild-type lobular breast cancer

Based on our identification of an inactivating *AFDN* frameshift mutation, we analyzed the occurrence and frequency of *AFDN* mutations in the publicly available breast cancer databases METABRIC and TCGA. Overall, in TCGA and METABRIC combined, 1/142 tumors (0.7%) have had a reported truncating mutation, which is similar to our study (1/146, 0.7%). In total, 79 unique somatic *AFDN* mutations were reported in a total of n=88 mammary carcinomas, of which 70 (79.5%) were IDC-NST (invasive ductal carcinoma, no special type), 10 (11.4%) ILC and 5 (5.6%) mixed Ductal and Lobular carcinoma cases (Suppl. Table 1). The remaining 3 cases (3.3%) were reported without specified histologic subtype. Among the 54 missense mutations, 34 (63.0%) were associated with decreased Afadin mRNA expression (Suppl. Table 1). Of the 27 *AFDN* truncating (Frame-Shift and Nonsense) mutations reported within METABRIC and TCGA, 24 mutations (88.9%) are predicted to cause loss of the C-terminal domain of *AFDN*, leading to loss of both the α-catenin binding site and FAB domain (Fig. 2A).

**Figure 2.**
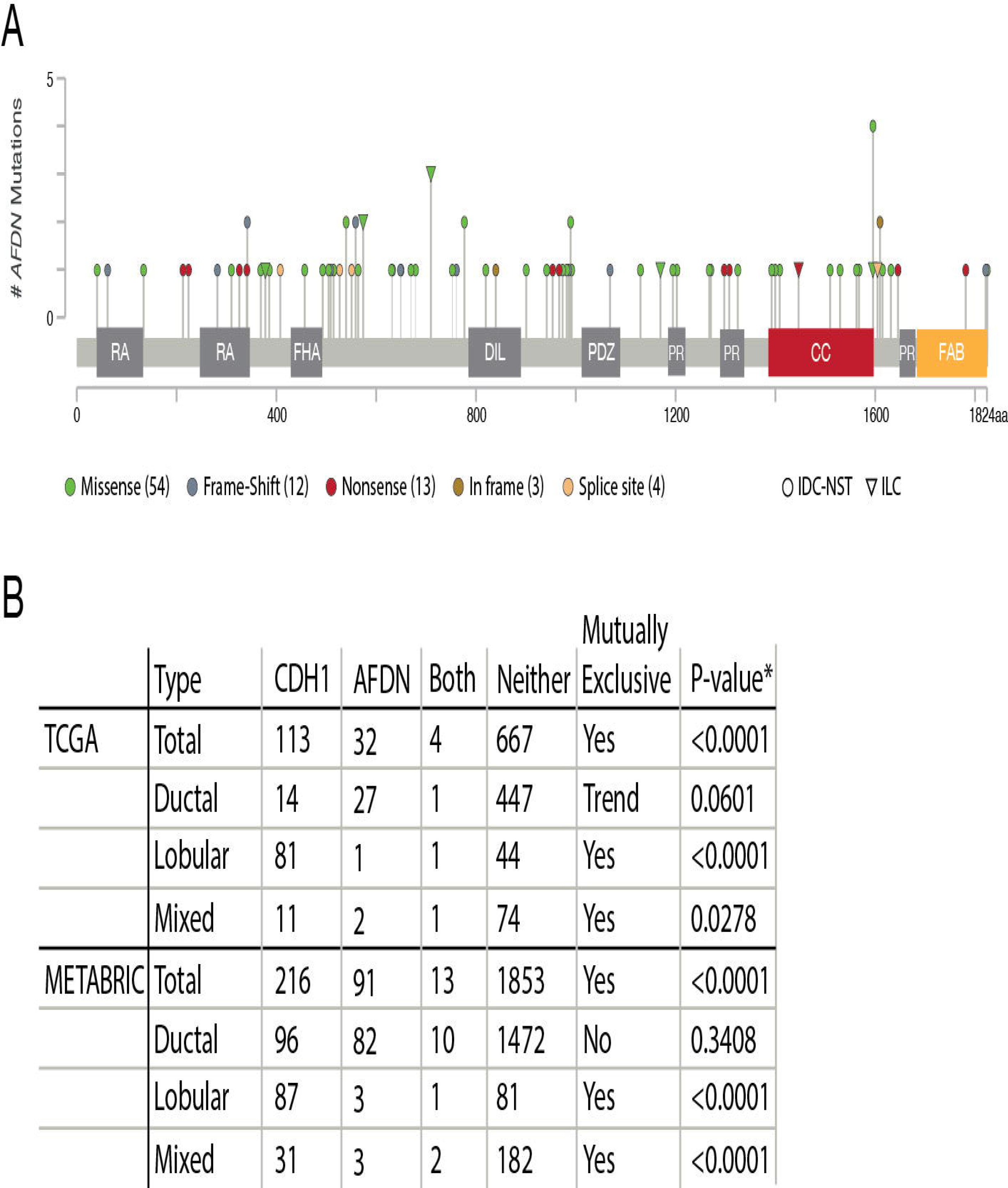
The occurrence of *AFDN* mutations in breast cancer predispose towards C-terminal protein loss and occur mutually exclusively from somatic *CDH1* mutations. A. METABRIC and TCGA cBioportal AFDN mutations (Pereira *et al*, 2016; Berger *et al*, 2018). Lollipop graph depicting distribution of *AFDN* mutations from the TCGA and METABRIC databases. Domains and regions: RA, Ras associated; FHA, forkhead associated; DIL, Dilution; PR, proline rich; CC, coiled coil; FAB, F-actin binding. B. TCGA and METABRIC mutual exclusivity of AFDN and CDH1 mutations in breast cancer. Somatic mutations in AFDN and CDH1 occur mutually exclusive in ILC and ILC-like breast cancer. * p-value calculated by Fisher’s Exact Test.

We next investigated if *CDH1* and *AFDN* mutations co-occurred in the METABRIC and TCGA breast cancer datasets and found significant overall mutual exclusivity for *CDH1* and *AFDN (*p<0.0001; Fig. 2B), especially in the subtypes ILC and mixed ductal/lobular carcinoma (p<0.0001; Fig. 2B). In contrast, and as expected, we found that *CDH1* and *AFDN* mutations do not occur mutually exclusive in IDC-NOS. Finally, we performed a blinded histopathological evaluation of 20 TCGA cases containing *AFDN* mutations to confirm the initial differential diagnosis. We revised one case that was initially diagnosed as IDC-NST as a mixed Ductal and Lobular phenotype (TCGA-BH-A0DZ-01, Suppl. Table 1). Of note, we revised two cases as apocrine breast cancer, a subtype that normally has an occurrence of 1% (Dellapasqua et al. 2013), compared to 10% in this subset of *AFDN* mutant tumors.

### Inactivation of *AFDN* leads to loss of cell-cell contact and anoikis resistance in breast cancer cells

To investigate the consequences of Afadin loss in breast cancer, we performed CRISPR/Cas9-mediated knockout of *AFDN* in MCF7 cells, an E-cadherin expressing and ER positive human breast cancer cell line (Fig. 3A). After transduction, Afadin knockout cells (MCF7::Δ*AFDN*) were selected, expanded and subjected to phenotypical analysis. In contrast to wild type MCF7 cells, MCF7::Δ*AFDN* cells did not show a cobblestone epithelial phenotype, but instead grew as non-cohesive motile cells (Fig. 3B). Because anchorage-independence is a hallmark of ILC cells (Derksen *et al*, 2011, 2006), we next assessed if Afadin loss induces anoikis resistance. Indeed, MCF7::Δ*AFDN* cells cultured in suspended (anchorage independent) conditions acquired anoikis resistance, comparable to E-cadherin loss in MCF7::Δ*CDH1* (Fig. 3C & D) (Hornsveld *et al*, 2016). Moreover, using inhibitors targeting RhoA and Rock1 (C3 and Y27632), we showed that the anoikis resistance of MCF7::Δ*AFDN* cells critically depends on Rho/Rock-dependent actomyosin contraction, comparable to the MCF7::Δ*CDH1* cells (Fig. 3E), allowing to conclude that Afadin loss induces an actomyosin-dependent anoikis resistant phenotype.

**Figure 3.**
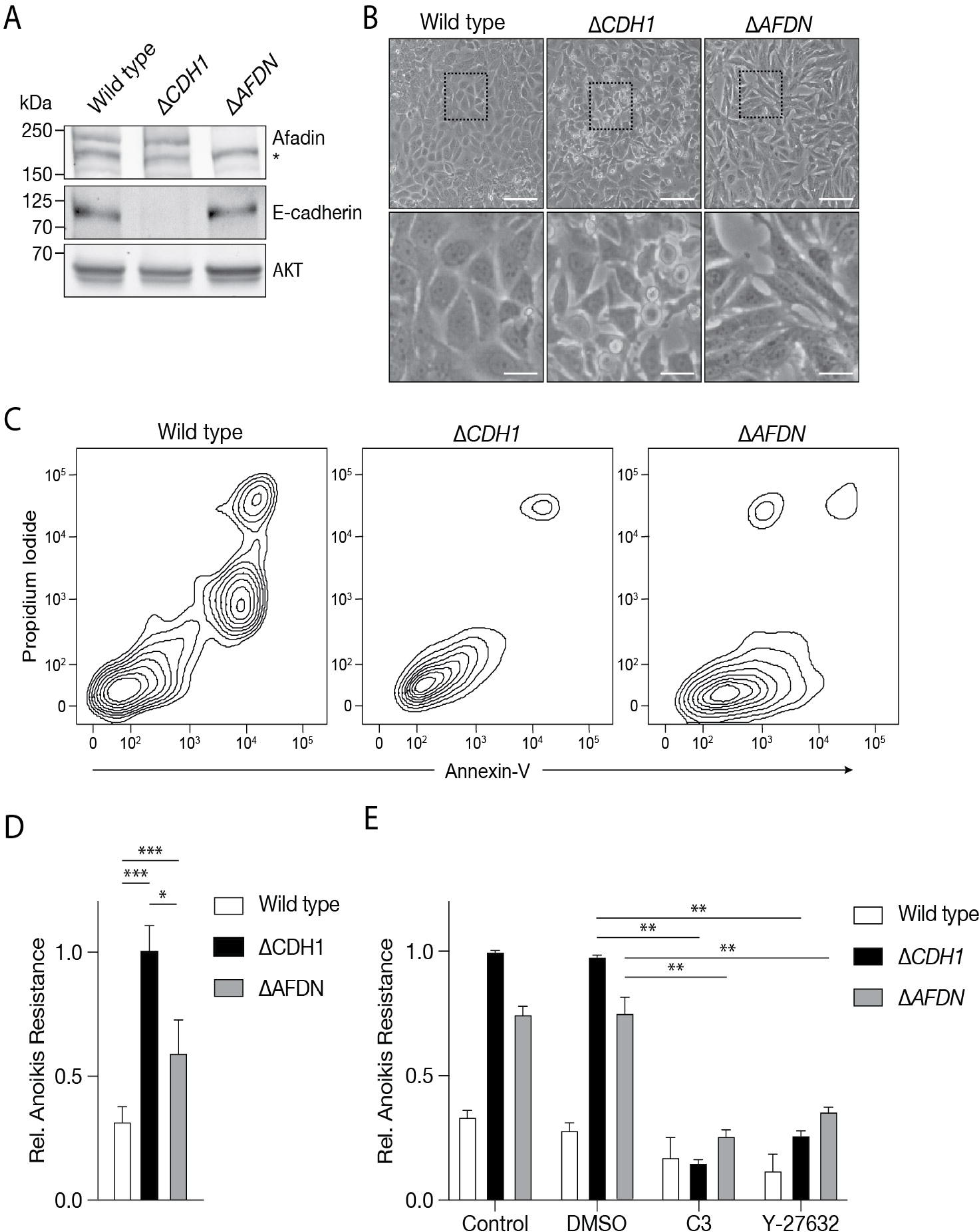
Loss of *AFDN* in E-cadherin expressing breast cancer cells MCF7 cells leads to loss of cell-cell adhesion and anoikis resistance. **A.** Western blot showing the effect of CRISPR-Cas9 mediated knockout of Afadin or E-cadherin in MCF7. AKT was used as loading control. *, unspecific protein (loading). **B.** Representative DIC images showing MCF7 wild type, MCF7::Δ*AFDN,* and MCF7::Δ*CDH1* cells, revealing a phenotypical change in cellular morphology upon *CDH1* or *AFDN* knockout. Bottom row shows zoom-ins of top row. Size bar = 100 µM top row, size bar = 20 µM bottom row. **C** and **D.** Afadin loss induces anoikis resistance. Contour plots (C) showing the anoikis resistance profiles of MCF7, MCF7::Δ*CDH1* and MCF7::Δ*AFDN* cells. Samples were stained with FITC-conjugated Annexin-V and Propidium Iodide and analyzed by FACS. Quantifications are shown in (D) **E.** Anoikis resistance in MCF7::Δ*AFDN* cells is RhoA and Rock1 dependent. Cells were cultured in suspended conditions and treated with C3 transferase (C3, Rho inhibitor) or Y-27632 (Rock inhibitor). Anoikis resistance was assessed as in (D). *p<0.05; **p<0.01 ***p<0.001.

### AFDN loss attenuates proper adherens junction formation and F-actin linkage

In contrast to MCF7 wild type cells, MCF7::Δ*AFDN* cells show an aberrant expression of E-cadherin on the plasma membrane, similar to the E-cadherin expression patterns in the *AFDN* mutant ILC sample. Immunofluorescence (IF) for Afadin confirmed the successful knockout (Fig. 4A). Concomitant IF for E-cadherin, β-catenin and α-catenin showed that an AJ was formed on the plasma membrane in MCF7::Δ*AFDN* cells (Fig. 4A, Suppl. Fig. 1A and 1B), but with a patchy, punctate expression pattern (Fig. 4A). In agreement with published literature (Sakakibara *et al*, 2020), we found that Zona Occludens 1 (ZO1) Fig. expression is reduced at the tri-cellular junctions compared to wild type controls (Suppl. Fig. 1C). Importantly, F-actin IF expression analysis showed that cortical F-actin distribution is less confined at the plasma membrane when compared to MCF wild type cells (Fig. 4A). To quantify these differences, we examined the spatial E-cadherin/F-actin expression and distribution at the plasma membrane using perpendicular line scans (Fig. 4B), and calculated the ratio of F-actin over E-cadherin expression at the Full Width at Half Maximum Height (FWHM). These analyses confirmed that loss of *AFDN* in MCF7::Δ*AFDN* cells leads to diffused distribution of cortical F-actin (Fig. 4C & D). Our findings therefore indicate that Afadin controls proper linkage from E-cadherin to the F-actin cytoskeleton. Furthermore, our work suggests that Afadin loss in breast cancer cells induces improper F-actin mechanical tension, leading to constitutive deregulation of actomyosin contraction, which has been causally linked to anoikis resistance in ILC (Schackmann *et al*, 2011; Groot *et al*, 2018).

**Figure 4.**
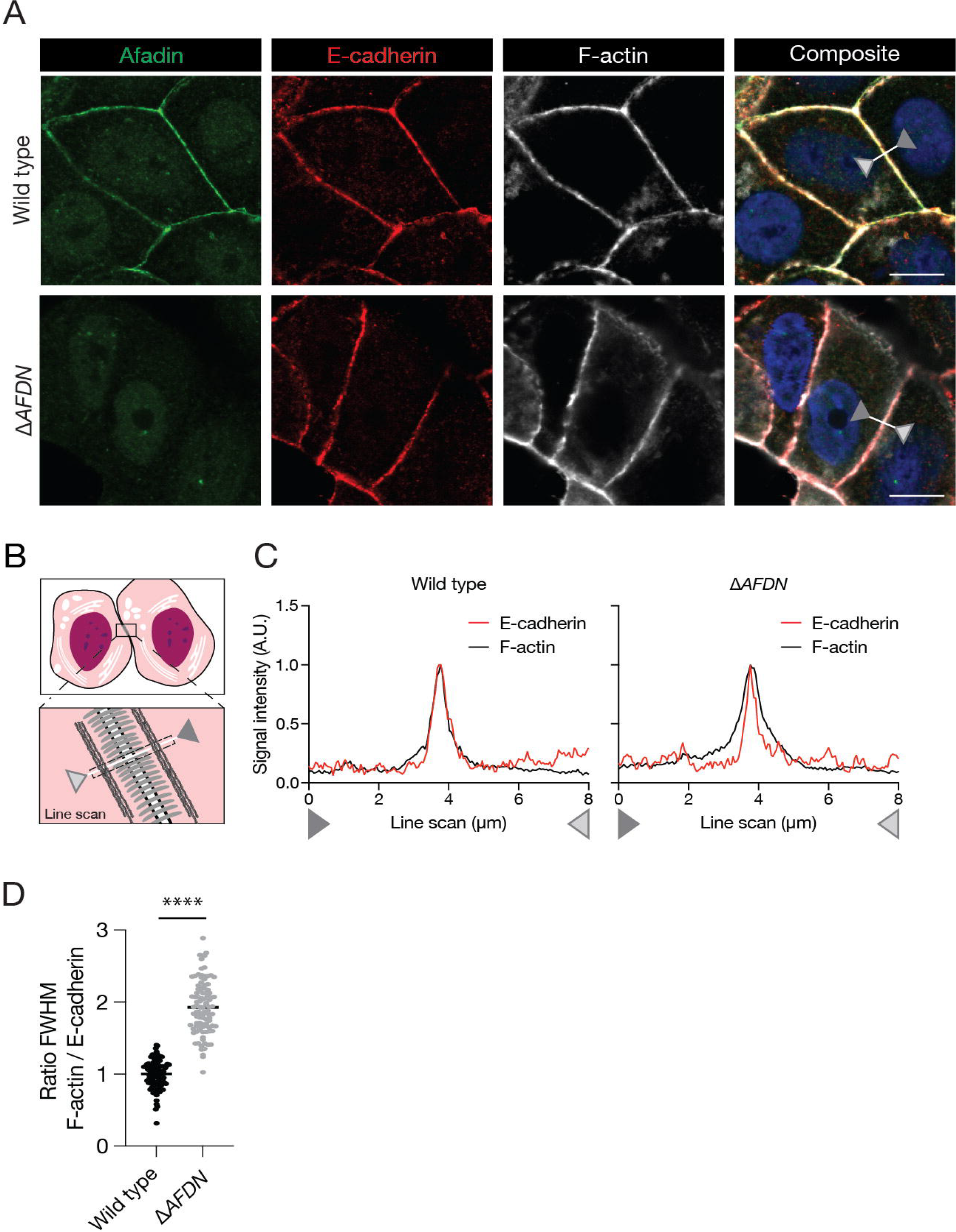
Afadin loss attenuates proper AJ formation and cortical F-actin distribution. **A.** Immunofluorescence for Afadin (green), E-cadherin (red) and F-actin (white), showing the attenuation of proper AJ formation. The right image is a composite of the left panels. Size bar = 5 µm. **B – D.** Requirement of Afadin for proper F-actin organization at the epithelial AJ. Cartoon (B) showing the setup of the line scan quantifications shown in (C and D). Line scans comparing the localization of E-cadherin and F-actin were done and quantified perpendicular to the plasma membrane (between the arrows indicated in the composite image depicted in (A)). Quantification of colocalization was done based on the ratio of the Full Width at Half Maximum (FWHM) of F-actin over E-cadherin from wild type MCF7 (black) *versus* MCF7::Δ*AFDN* cells. ****p<0.0001.

### The CC domain of Afadin is essential for proper AJ formation

The identified *AFDN* mutations indicate that most truncating mutations lead to loss of the C-terminal domain, causing impaiment of binding to α-catenin and F-actin respectively. We therefore hypothesized that an Afadin dependent F-actin to AJ connection occurs through either binding of α-catenin to the Afadin CC-domain, through the Afadin FAB-domain, or both. To investigate the individual requirement, we used full length N-terminally GFP-tagged rat l-Afadin (FL) (Sakakibara *et al*, 2018) and generated truncated constructs lacking either the CC-domain containing the α-catenin binding region (ΔCC) the FAB-domain (ΔFAB), or the entire C-terminus from amino acid position 1391 onward (ΔCterm)(Fig. 5A). We next performed reconstitution experiments in MCF7::Δ*AFDN* cells, confirming expression at the correct molecular weight of either the full length or truncated constructs using a GFP antibody (Fig. 5B). Expression of either the FL or ΔFAB constructs in MCF7::Δ*AFDN* cells fully rescued the non-cohesive phenotype, resulting in proper cell-cell adhesion and a cobblestone morphology (Fig. 5C). In contrast, reconstitution with either the ΔCC or ΔCterm truncates did not restore the epithelial phenotype in MCF7::Δ*AFDN* cells (Fig 5C). Although both truncates did not rescue epithelial morphology, cells reconstituted with ΔCC constructs showed a hypomorphic phenotype, *i.e.* a partial retainment of the migratory morphology and spindle-shaped cells (Fig. 5C). In line with the cellular phenotypes, subsequent examination and quantification of the spatial E-cadherin/F-actin expression at the plasma membrane confirmed that the FL and ΔFAB reconstitution fully rescue proper co-localization of cortical F-actin to E-cadherin (Fig. 5D). In contrast to the FL and ΔFAB rescue experiments, we observe that neither the ΔCC, nor the ΔCterm reconstitution is able to rescue Afadin loss, leading to higher F-actin/E-cadherin ratios, *i.e.* dispersed cortical F-actin (Fig. 5E). Our reconstitution experiments in AFDN knockout cells thus show that the coiled-coil region of Afadin that contains the α-catenin binding domain, is required for proper AJ formation and linkage to the F-actin cytoskeleton in E-cadherin expressing breast cancer cells. We conclude that attenuated cortical F-actin distribution may underpin the acquisition of Rho-Rock-dependent anoikis resistance upon Afadin inactivation.

**Figure 5.**
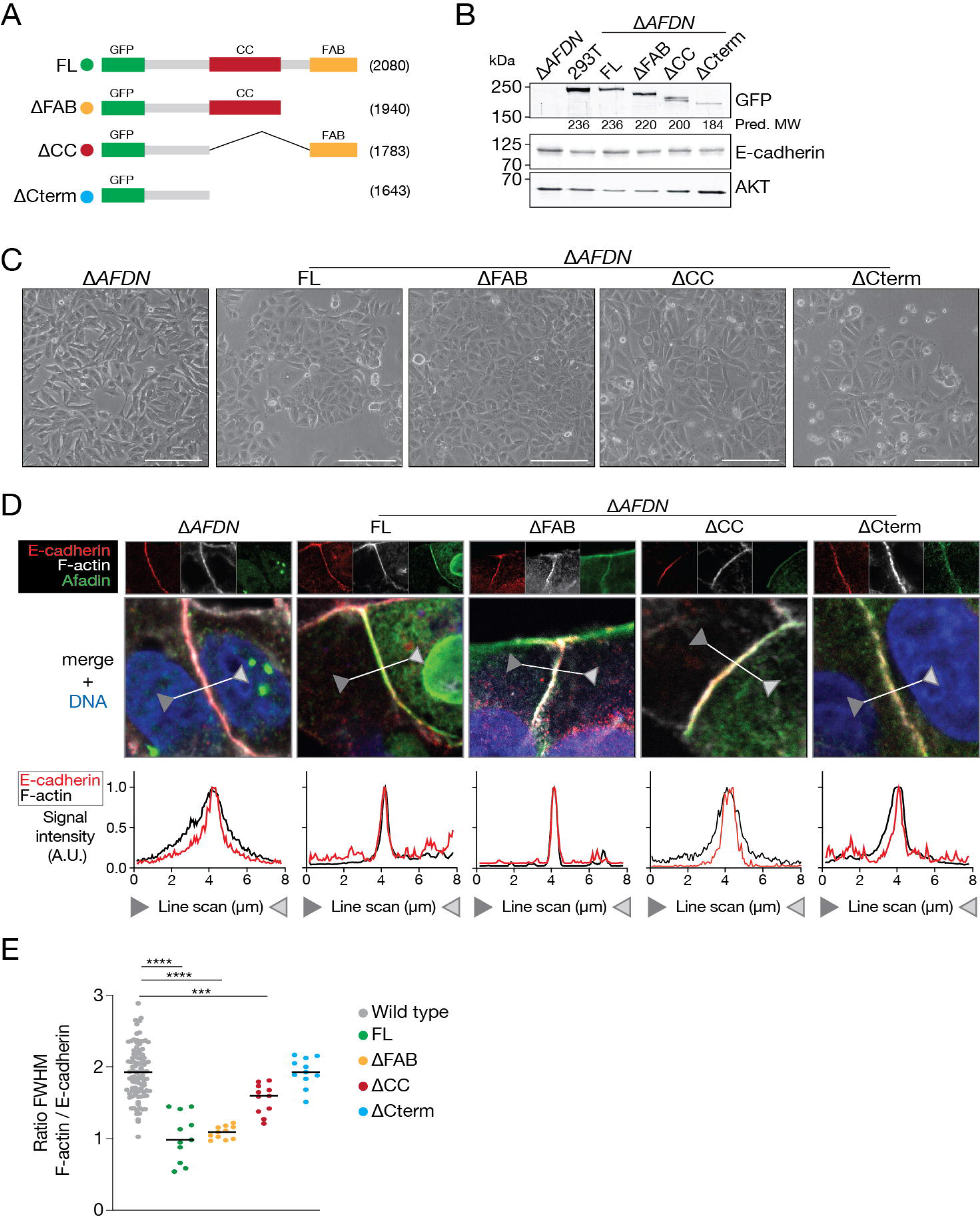
The α-catenin binding domain of Afadin is essential for epithelial cell-cell adhesion and correct cortical F-actin localization. **A.** Schematic overview of the GFP-tagged full length and truncation Afadin. **B** and **C.** Reconstitution of Afadin full length or truncates in MCF7::Δ*AFDN* cells. Western blots (B) showing expression based on the GFP tag (top, GFP). The impact of Afadin loss or truncation on E-cadherin expression was probed (middle), and Akt was used as loading control (bottom). The predicted molecular protein weight is indicated below the GFP blot (Pred. MW). HEK293Ts were used as a transfection control with the FL construct. Representative DIC images (C) showing the phenotypical impact of the Afadin truncates in MCF7::Δ*AFDN* cells. Similar cell numbers were plated and photographed after 18 hours. Size bar = 10 µm. **D** and **E.** Immunofluorescence images showing MCF7::Δ*AFDN* cells reconstituted with the indicated truncates (see Fig. 5A for details). E-cadherin, F-actin and GFP-Afadin were stained (D) and line scans were performed between the arrow heads (indicated in the merged composites). The ratio of the full width at half maximum (FWHM) of the F-actin over E-cadherin expression signals were quantified, normalized to 1, and compared in (E). ***p<0.001; ****p<0.0001. Note that absence of the Afadin F-actin binding domain is not sufficient to disrupt E-cadherin to F-actin linkage, while deletion of the α-catenin domain is required for proper AJ formation.

### Loss of *AFDN* leads to an ILC phenotypical features and lung metastasis phenotype in a transplantation-based mouse model of breast cancer progression

To assess the functional impact of *AFDN* loss in breast cancer, we performed mammary intraductal injections, using MCF7::Δ*AFDN* cells that had been stably reconstituted with either the ΔCC or ΔCterm GFP-tagged AFDN truncate. In this setup we monitored longitudinal tumor volumes and observed no significant differences in primary tumor size between the tumor cohorts, indicating that Afadin loss does not impact primary tumor growth (Fig. 6A). All cohorts had retained membranous expression of E-cadherin (Fig. 6B, middle panels) with carcinomas in the rescue cohorts expressing GFP, confirming proper expression of the constructs after transplantation (Fig. 6B, bottom panels). Moreover, carcinomas in the wild type MCF7 tumor cohort were characterized by expansive, circumscribed growth with occasional collective invasion of cells into the duct (Fig. 6C). In contrast, mice injected with MCF7::Δ*AFDN* or the ΔαCAT/ΔCterm reconstituted cells showed typical ILC-like features, including overt single cell invasion and the formation of trabeculae and single or Indian files, most notably at the invasive front (Fig. 6C). Interestingly, we observed a significantly shorter overall survival of mice that were injected with either the MCF7::Δ*AFDN* knockout cells or the ΔCterm truncate reconstituted cells (p<0.05)(Fig. 6D). This was supported by single cell intravasation of cancer cells in the MCF7::Δ*AFDN* knockout cells and both the ΔCC and ΔCterm reconstitution conditions (Fig. 6E). Examination of the lung tissues showed that the decreased survival in these mice was due to the development of overt metastasis of cells expressing human GATA3, used as a marker for human cells (Fig. 6F and 6G). Although reconstitution with ΔCC did not result in a significant survival decrease, we detected an increase in metastatic burden after quantification of the GATA3 positive cells, similar to the increase observed after reconstitution with the ΔCterm truncate (Fig. 6F and 6).

**Figure 6.**
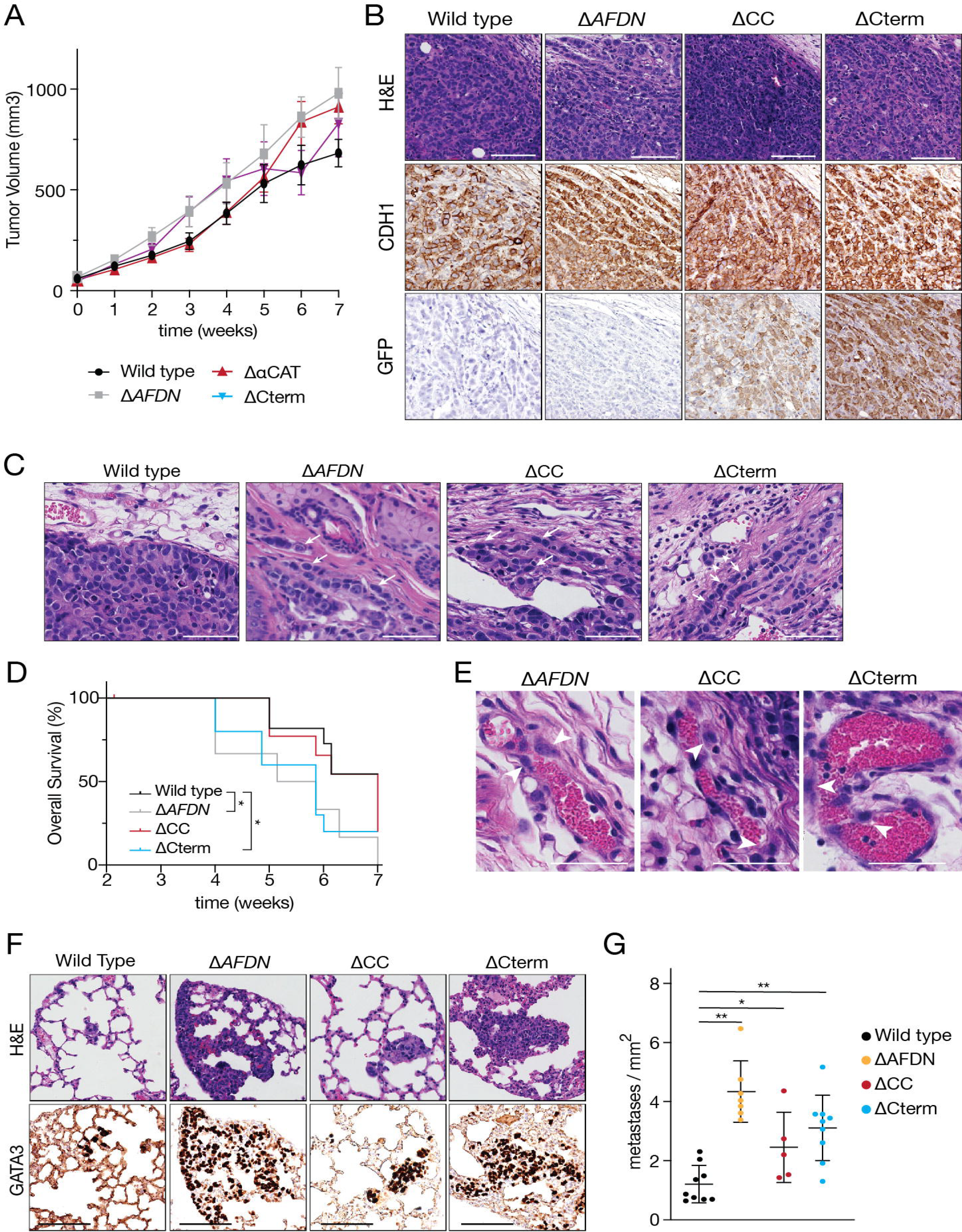
Loss of Afadin leads to metastatic breast cancer with invasive lobular features in mice. **A.** Inhibition of proper AJ formation does not impact tumor growth The indicated MCF cells were injected intraductally in mice, and tumor volumes quantified over time. Error bars depict standard deviations. **B.** E-cadherin expression is retained in cells lacking Afadin or Afadin C-terminal truncates. Shown are immunohistochemistry panels for E-cadherin (middle) and GFP-Afadin (bottom) in the indicated primary tumors. Left panels are H&E stains. Size bar = 100 µm. **C.** Afadin loss of function induces ILC features. Shown are representative examples of primary tumors that developed in the mice from (A). Note the acquisition of invasive trabecular single cell files (arrows) and individual cells invading (arrowheads). Size bar = 100µm. **D.** Loss of the Afadin C-terminal α-catenin and F-actin binding domains results in reduced survival. Shown is a Kaplan Meijer tumor-free survival plot for the indicated tumor cohorts. Censored cases are marked by ticks on the plots. **E.** Examples of intravasation in the periphery of primary tumors that developed in the mice from (A). Arrowheads indicate individual tumor cells. Size bar = 50µm. **F and G.** Loss of the Afadin C-terminal α-catenin is sufficient to drive metastatic disease. Shown are H&E stains (top) and immunohistochemistry for human GATA3 (bottom) on lung sections (D). Note the overt increase in metastatic potential upon loss of the α-catenin binding domain of Afadin. Size bar = 50 µm. **G.** Quantification of metastases, normalized to lung area.

We conclude that loss of Afadin expression in E-cadherin expressing breast cancer cells induces the formation of ILC-type metastatic tumors. Our data reveal an essential tumor suppressive role for Afadin in breast cancer progression. Afadin is essential to control a proper and matured connection of E-cadherin to the F-actin cytoskeleton to prevent loss of cell-cell contacts, the acquisition of anoikis resistance, and the progression towards ILC-like metastatic tumors (see Fig. 7). In sum, we have identified Afadin as a candidate tumor suppressor for ILC progression.

**Figure 7.**
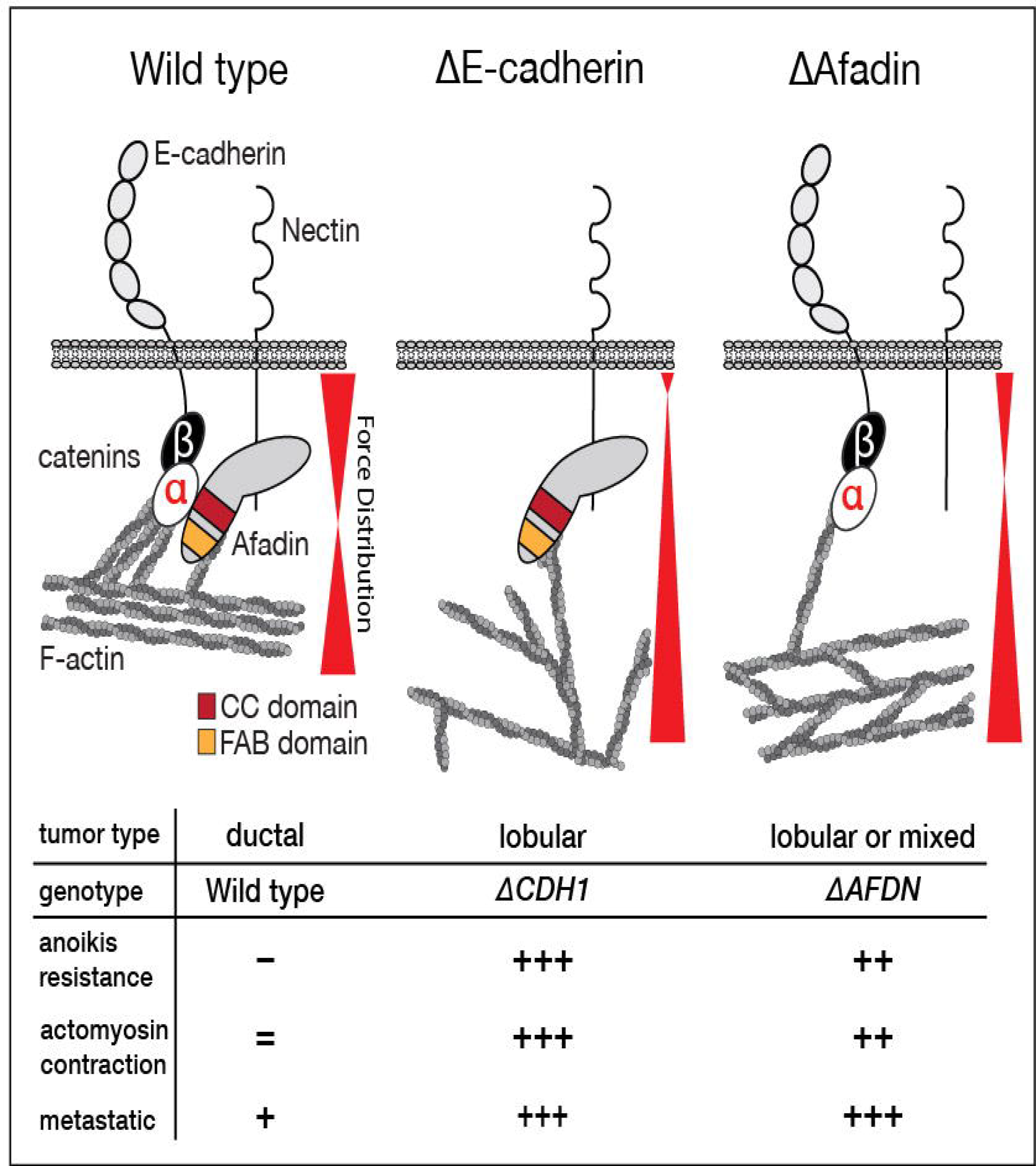
Working model for the consequences of Afadin loss on the interaction of the Adherens Junction and the F-actin cytoskeleton. In *CDH1* wild type conditions, the AJ can form proper interactions with the F-actin cytoskeleton, leading to a balanced outside-in force distribution from the AJ to the F-actin cytoskeleton and complete maturation of the AJ. Tumors with an intact and balanced AJs will not acquire full anoikis resistance and display low metastatic potential (left panel, Wild type). Upon mutational loss of E-cadherin, cells are unable to form E-cadherin based AJs, leading to overt loss of mechanical forces from the AJ to the F-actin cytoskeleton due to attenuated feed-back signals (middle panel, ΔE-cadherin). E-cadherin loss results in constitutive actomyosin contraction, Rho-Rock dependent anoikis resistance, and characteristic ILC-type phenotypes like trabecular growth patterns, single cell invasion, and distant metastasis. Upon Afadin loss, E-cadherin based-AJ are formed, but the lack of stabilization that occurs from inability of Afadin to bind the α- catenin domain containing coiled-coil region, causes a disbalance of forces towards the F-actin cytoskeleton, which prevents proper formation of a mature AJ (right panel, ΔAfadin). The classical C-terminal F-actin binding domain is dispensable for this function. As a result of unbalanced mechanical forces, adequate feedback from the F-actin cytoskeleton is prevented, and homotypic E-cadherin based cell-cell junction cannot mature properly, resulting in dispersed cortical F-actin distribution. Consequently, cells display and depend on actomyosin contraction driven anoikis resistance, leading to invasion and single cell breast cancer metastasis.

## DISCUSSION

Loss of E-cadherin underpins ILC biology and etiology (reviewed in: (Christgen *et al*, 2016)). The biochemical consequences of E-cadherin loss are converging on the activation of survival pathways driven by growth factor receptor activation (Bajrami *et al*, 2018; Nagle *et al*, 2018; Teo *et al*, 2018) and constitutive actomyosin contraction (Schackmann *et al*, 2011; Ven *et al*, 2015; Groot *et al*, 2018). Complete loss of E-cadherin expression in ILC appears to drive actomyosin contraction through loss of cell contacts and autocrine activation of RhoA through the suppression of the Rock antagonist MRIP (Schackmann *et al*, 2011). However, it remained less clear what propels F-actin contraction in the 10% of ILC cases that have retained E-cadherin expression.

Previous data have connected loss of α-catenin to E-cadherin expressing ILC cell lines (Hazan *et al*, 1997; Hollestelle *et al*, 2010), and provided functional evidence that loss of α-catenin leads to an ILC-like phenotype and actomyosin-dependent anoikis resistance (Groot *et al*, 2018). In short, the aforementioned studies indicated that an imbalanced actomyosin-dependent force distribution due to loss of proper linkage of E-cadherin to the F-actin cytoskeleton, may be sufficient to drive single cell survival and dissemination during ILC progression. In the current study we have used a combination of genomics, cell biology and mouse experiments to identify new alternative drivers of ILC. Based on the strong dependency on a balanced E-cadherin to F-actin connection, we have performed targeted next generation DNA sequencing on ILC cases that retained the wild type *CDH1* alleles and identified the *AFDN p*.Lys630stop frameshift mutation. Afadin is linked to the metastasis free survival of breast cancer (Letessier *et al*, 2007; Fournier *et al*, 2011), and recent retrospective phylogenetic analyses identified a *CDH1* wild type ILC case that presented with an *AFDN* deletion (Fimereli *et al*, 2022). Interestingly, we found that 82% of the identified *AFDN* mutations in the TCGA and METABRIC databases (0.4% of total) occur in IDC-NST, which suggests that some cases were potentially misdiagnosed and may in fact histologically be ILC or mixed IDC/ILC. Indeed, from the twenty *AFDN* mutant TCGA cases that had available histology data, we could identify one case being misdiagnosed as IDC-NST.

Experimentally, loss of Afadin in E-cadherin expressing luminal-type MCF7 cells leads to less confined cortical localization of F-actin at the plasma membrane and subsequent colocalization with the E-cadherin based AJ, indicative of incomplete maturation of the AJ. With the residual but aberrant localization of wild type E-cadherin complexes on the membrane, we conclude that the tumor suppressor function of Afadin is derived from the attenuated association and incomplete maturation of the E-cadherin-F-actin cytoskeleton link. In the case of Afadin loss, we found that the coiled-coil domain containing an α-catenin binding region is necessary and sufficient to confer proper actomyosin organization at the AJ. Because decreased cortical F-actin localization and contraction are caused by reduced buildup of forces along the F-actin fibers connecting the cortical F-actin to the AJ (Vasioukhin *et al*, 2000; Verma *et al*, 2004; Sumida *et al*, 2011; Taguchi *et al*, 2011), we conclude that this lack of pulling forces is strongly exacerbated in the absence of Afadin, where insufficient coupling of E-cadherin to F-actin prevents apical constriction. Consequently, Rho Kinase may be released from Shroom3 and drive actomyosin contraction (Kimura *et al*, 1996; Uehata *et al*, 1997), a process that will eventually drive ILC formation and metastasis. One of the causes of the reduction in mechano-transduction upon Afadin loss could be the forces of F-actin exerted on Vinculin during initial AJ formation, which is essential for establishing the AJ and F-actin connection (Duc *et al*, 2010). It has become clear that α-catenin is the rate-limiting tension sensor upon stretch activation between E-cadherin and F-actin (Yonemura *et al*, 2010), whereby Vinculin interacts with α-catenin only after unfolding of α-catenin (Yao *et al*, 2014). Upon AJ maturation following a balanced interplay of forces, localized expression levels of Vinculin are reduced in the junction, rendering a stabilized AJ including Afadin (Rooij, 2014). Previous work by the Takai laboratory demonstrated that Afadin is required for proper AJ formation, but that the FAB domain is dispensable for this event (Sakakibara *et al*, 2020). Our work is in full agreement with these findings, showing that reconstitution of an Afadin truncate lacking the C-terminal FAB domain fully rescues correct formation of the E-cadherin to F-actin connection, and the restoration of an epithelial phenotype. In contrast, we show that reconstitution with deletion truncates lacking the CC domain prevents cortical actin colocalization with the E-cadherin based AJ, which shows that direct association of the CC domain and Afadin upon AJ formation is key in establishing mechano-responsive interactions to the F-actin cytoskeleton.

It is nonetheless striking that the C-terminal FAB domain is dispensable for proper AJ formation and function in breast cancer cells. Previous studies in Drosophila have demonstrated that while the FAB domain is not essential for AJ formation, it supports the stability of AJ under tension (Perez-Vale *et al*, 2021). In non-malignant mammalian cells, loss the FAB domain appears to delay the formation of adherens and tight junctions (Sakakibara *et al*, 2018). Interestingly, the CC domain of Afadin contains four predicted alpha helices, of which the first N-terminal helix overlaps with the mapped binding site of Afadin to α-catenin (Maruo *et al*, 2018). Recent analyses by the Peifer lab indicate that the predicted C-terminal helices show a near complete overlap with the F-actin binding site identified by Carminati *et al*. (Carminati *et al*, 2016), suggesting that the CC domain might not only confer α- catenin binding, but also F-actin interaction, or both (preprint Gurley *et al*, 2023). In short, based on our work and the aforementioned studies we propose that either: (i) the FAB domain binds F-actin and enforces AJ stability but is not solely responsible for F-actin linkage, (ii) all major functions related to AJ stability are deployed by Afadin through α-catenin dependent F-actin linkage, or (iii) the predicted F-actin binding sites in the CC intrinsically disordered region (IDR) are dominant in facilitating AJ formation and stability together with α-catenin linkage. Because we observe that deletion of the CC domain is sufficient to disturb epithelial integrity in breast cancer cells through improper AJ formation, induce actomyosin dependent anoikis resistance, and drive invasive tumor growth and metastatic dissemination, it remains currently unclear how interplay of the scenarios controls the phenotypes observed. Future work will have to delineate the exact mechanisms that underpin the role of Afadin in AJ formation and function through fine-mapping of the predicted F-actin sites with the IDR of Afadin.

Regardless of the precise mode of action, we envision that in parallel to the Afadin-α-catenin bond, the concomitant linkage of Afadin to F-actin serves to further stabilize the AJ and allow subsequent build-up of forces that can then reach a threshold for the reduction of Vinculin levels. In this scenario, Afadin would function as a gatekeeper of AJ maturation through the exposure of force dependent unfolding of the Afadin binding MIII subdomain/region of α-catenin (Sakakibara *et al*, 2020). While the exact balance of forces and domain requirements are still unclear, our work indicates that the necessity of Afadin for proper maturation of the epithelial AJ acts as a potent tumor suppressor mechanism for the development and progression of ILC. Ultimately, preventing formation of a mature AJ and the accompanying mechanical transduction may lead to constitutive activation of RhoA and Rock1 dependent actomyosin contraction through the disturbance of physiological feedback. Our data indicate that these events lead to anoikis resistance, a feature that underpins metastatic dissemination (Douma *et al*, 2004; Derksen *et al*, 2006).

In conclusion, loss of *AFDN* leads to an ILC-like phenotype in E-cadherin expressing breast cancer cells, due to its binding to α-catenin and the subsequent linkage to, and stabilization of, the F-actin cytoskeleton. Similar to the consequence of α-catenin inactivation in breast cancer cells (Groot *et al*, 2018), we demonstrate that Afadin loss leads to an F-actin dependent non-cohesive cellular phenotype, founded in detrimental cortical actomyosin dynamics that result in anchorage independence and metastatic dissemination. As such, we reaffirm the notion that carcinomas driven by loss of the AJ or the loss of the F-actin cytoskeleton connection with the AJ should be considered ‘actin’ driven diseases (Bruner & Derksen, 2018). Delineating the disruption of proper AJ formation will be key in the correct diagnosis and future treatment of malignancies driven by functional loss of E-cadherin such as ILC and diffuse gastric cancer.

## SUPPLEMENTAL FIGURE LEGENDS

**Supplemental Figure 1. A-C** Immunofluorescence images showing single and merged images (right panels) of E-cadherin and α-catenin (A), E-cadherin and β-catenin (B) and Afadin and ZO-1 (C) in MCF7 wild type (Wild type) and MCF7::Δ*AFDN* cells (Δ*AFDN*). Note the loss of ZO-1 expression at the tri-cellular junctions. Size bar = 5 µm.

**Supplemental Figure 2.** Afadin function does not impact Estrogen receptor (ER) expression in primary tumors. Shown are H&E (left panels) and ER expression (right panels) in primary tumors that developed after injection with either MCF7 wild type, MCF7::Δ*AFDN*, or MCF7::Δ*AFDN* cells, expressing either the ΔCC or ΔCterm Afadin truncates.

## SUPPLEMENTAL TABLE HEADINGS

**Supplemental Table 1. TCGA & METABRIC *AFDN* status.** Full overview of all *AFDN* mutations reported in TCGA and METABRIC with revised diagnoses and *CDH1* status.

**Supplemental Table 2.** gRNAs and primers used for the CRISPR/Cas9 mediated knockout of *AFDN* and subsequent TIDE analysis and primers designed to generate the Afadin Full Length, ΔFAB, ΔCC and ΔCterm truncate constructs via In-Fusion PCR.

## AUTHORSHIP CONTRIBUTIONS

M.A.K.R. and P.W.B.D. designed the study. M.A.K.R., N.F.L.E, and K.V performed experiments. N.F.L.E. and M.A.K.R. performed preclinical mouse experiments. P.W.B.D., I.N. and J.L. designed the targeted sequencing capture strategy and performed subsequent analyses. R.B. provided patient data, tissue samples and DNA samples. N.I. provided conceptual input in Afadin form, function, and cloning of Afadin truncate constructs. M.C. and P.D. performed human and mouse histological tumor diagnostics. All authors read the manuscript and were given opportunity to provide input.

## ACKNOWLEDGEMENTS

We acknowledge the Utrecht Sequencing Facility (USEQ) for providing sequencing service and data. USEQ is subsidized by the University Medical Center Utrecht and The Netherlands X-omics Initiative (NWO project 184.034.019). We thank Astrid Bosma for help with patient samples and Yoshimi Takai for the full length rat I-Afadin construct. Financial support was obtained from the Dutch Cancer Society (KWF 10456), Breast Cancer Now (2018NovPCC1297), and the European Union’s Horizon 2020 FET Proactive program under the grant agreement No. 731957 (MECHANO-CONTROL). This publication is also based upon work from COST action LOBSTERPOT (CA19138), supported by COST (European Cooperation in Science and Technology) https://www.cost.eu.

## MATERIALS AND METHODS

### Patient Material and Adhesome Sequencing

DNA extracted from ILC samples was kindly provided by the RATHER consortium (Michaut *et al*, 2016). Processed DNA was captured using a custom Agilent SurePrint array as previously described, based on the Adhesome as listed in Table 1. Sequencing of enriched samples was performed on an Illumina Miseq sequencer to a depth of >50x average target base coverage. All sequencing data were analyzed for mutations using in-house developed and validated pipelines (Mokry *et al*, 2010; Nijman *et al*, 2010; Harakalova *et al*, 2011; Hoogwater *et al*, 2010). All mutations were mapped to GRCh37.

### Cell Culture

MCF7 cells were cultured in DMEM-F12 (Invitrogen) containing 12% fetal calf serum, 100 IU/mL penicillin, and 100 μg/mL streptomycin. MCF7 was obtained from American Type Culture Collection and profiled using short-tandem repeat (STR) profiling (LGC Standards).

### CRISPR/Cas9-generated *AFDN* KO Baculovirus

Baculoviruses expressing Cas9 were produced as described previously (Hindriksen *et al*, 2017). Two gRNAs were tested and after TIDE analysis the highest efficiency gRNA was expanded (Brinkman *et al*, 2014). gRNAs and primers used to perform TIDE analysis are listed in Supplemental Table 2.

### Anoikis assay

Anoikis resistance was quantified as described previously (Schackmann *et al*, 2013). Samples were treated with: 10 µM Y-27632 (Selleckchem, Huissen, The Netherlands) and 0.02 µg/mL C3- transferase inhibitor (CT04-A cytoskeleton Inc, Heerhugowaard, The Netherlands), co-cultured with cells over 4 days in anchorage-independent conditions using ultra-low cluster 24 well plates (Corning). Error bars depict standard deviation. A two-sided Students’ t-test was performed to calculate statistical significance; p-values less than 0.05 were considered significant.

### Immunofluorescence

Cells were allowed to adhere to glass coverslips overnight, washed with PBS containing Ca^2+^ and Mg^2+^ (serial 14080055, Gibco, Paisley, Scotland, UK) and subsequently fixed using 4% paraformaldehyde. Samples were blocked with 4% BSA in PBS for 1 hr. Samples were incubated with primary antibodies overnight at 4°C. The following primary antibodies were used: mouse anti-p120 6H11 (1:500; Santa Cruz Biotechnology, Santa Cruz, California, United States), rabbit anti-GFP (1:1000; FL Santa Cruz), rabbit anti-l-Afadin (1:500, Sigma-Aldrich, Zwijndrecht, the Netherlands), monoclonal rat anti-E-cadherin (1:300, Sigma-Aldrich), mouse anti-p120-catenin (1:400, BD Biosciences, Becton, United States, clone98/pp120), mouse anti-α-catenin(1:100, Enzo Life Sciences, Farmingdale, New York, United States). For detection, samples were incubated with secondary antibodies in blocking buffer for 1 hr. at room temperature. The following secondary antibodies were used: (1:600; Invitrogen, Waltham, Massachusetts, United States): goat anti-mouse Alexa488 (#A11029), goat anti-rabbit Alexa488 (#A11034), goat anti-mouse Alexa568 (#A11031), goat anti-rabbit Alexa568 (#A11036), goat anti-rabbit Alexa647 (#A21245) and goat anti-rat Alexa647 (#A21247). Afterwards, cells were incubated with Alexa633-phalloidin to visualize F-actin (1:300; #A22284; Life Technologies, Carlsbad, California, United States) or DAPI to stain DNA, washed and mounted using Prolong Diamond Antifade Mountant (P36961; Thermo Fisher Scientific, Waltham, Massachusetts, United States). Imaging was performed on a LSM700 confocal microscope (Zeiss, Jena, Germany). Image processing was performed using ImageJ, Illustrator and Photoshop (Adobe).

### DNA Constructs

pEGFP-l-Afadin containing the full rat Afadin cDNA was a kind gift from Yoshimi Takai (Sakakibara *et al*, 2018). N-terminally GFP conjugated Afadin truncates were cloned into a pLV lentiviral backbone (pLV.bc.PURO, (Schackmann *et al*, 2011)) by In-Fusion PCR cloning using the pEGFP-l-Afadin plasmid as template. In short, pLV.BC.PURO was linearized using *Mlu*I and *Nhe*I digestion and the amplified GFP-Afadin fragments were incorporated using In-Fusion HD Cloning Plus (#638920; Takara, Kusatsu, Shiga, Japan). In-Fusion compatible complementary ends to the target vector were incorporated in the primers depicted in Supplementary Table 2.

### Transfections

Cells were plated at a density of 1*10^6 per 10cm plate and transfected on the following day using Fugene HD (Promega, Madison, Wisconsin, United States, E2311). A PGK-GFP vector was transfected in parallel as control for transfection efficiency. Three days after transfection, cells were selected for two days using 2 µg/mL Puromycin (Sigma Aldrich, P8833).

### Intraductal injections and mouse studies

MCF7::Δ*AFDN* cells were transduced with lentiviral pLV.BC.PURO particles with the different truncates as described (Schackmann *et al*, 2011). Two days after transduction, cells were put on Puromycin selection (2 µg/mL) for 2 weeks and expression was confirmed on Western Blot. 250,000 cells suspended in 25 µL were injected intraductally in the fourth mammary gland of recipient Rag2^-/-^;IL2cγR^-/-^ mice ((Gimeno *et al*, 2004), Envigo, Indianapolis, Indiana, United States) Tumor onset and growth was monitored longitudinally and tumor volume was determined by digital pressure-sensitive caliper measurements (Mitutoyo, Veenendaal, The Netherlands) using the following formula: 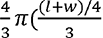). Animals that developed tumors were euthanized if the tumors reached a size of > 1000 mm^3^ or in cases of severe discomfort otherwise.

### Histological analysis

Formaldehyde-fixed, paraffin-embedded tissues were sectioned at 4 μm and stained with hematoxylin and eosin (H&E). For immunohistochemical staining, fixed sections were rehydrated and incubated with primary antibodies against ER (SP1; 1:1; Roche, Basel, Swiss), GATA3 (L50-823; 1:1; Roche), E-cadherin (612130; 1:150; BD Biosciences), and GFP (SC-8334;1:150; Santa Cruz). Endogenous peroxidases were blocked with 3% H2O2and biotin-conjugated secondary antibodies were used, followed by incubation with HRP-conjugated streptavidin–biotin complex (DAKO). Substrate was developed with DAB (DAKO)and pictures were produced using a Nikon Eclipse E800microscope with a Nikon DXM1200 digital camera (Nikon, Amsterdam, The Netherlands). Appropriate positive (expressing tissues) and negative controls (matched isotype and omitting primary antibody) were used throughout.

### Biochemistry

Cells were lysed in sample buffer and proteins were separated and western blotted as described previously (Schackmann *et al*, 2013). The following primary antibodies were used: Rabbit anti-l-Afadin (1:1.000, A0349; Sigma-Aldrich), rabbit anti-GFP (1:500; sc-8334, Santa Cruz Biotechnology), and goat anti-AKT (1:1,000, C-20/sc-1618; Santa Cruz Biotechnology). All blots were incubated for 30 min. with either rabbit anti-goat-PO (DAKO, Heverlee, Belgium; P160), goat anti-rabbit-PO (Bio-Rad, Veenendaal, The Netherlands; 170-6515) or goat anti-mouse-PO (Bio-Rad; 170-6516) secondary anti-bodies. Images were acquired on an Amersham Imager 600 (Amersham, ‘s-Hertogenbosch, The Netherlands).

### cBioportal analysis and data extraction

Data for the METABRIC and TCGA datasets were retrieved from the cBioportal public databases (https://www.cbioportal.org/) (Pereira *et al*, 2016; Cerami *et al*, 2012; Gao *et al*, 2013)

### Statistical Analysis

All statistical tests were performed using Prism 9 Version 9.4.1.

