## Supplementary figures and images for "Somatic mutation of Afadin leads to anchorage independent survival and metastatic growth of breast cancer through αE-catenin dependent destabilization of the adherens junction"

### Supplemental Figure 1

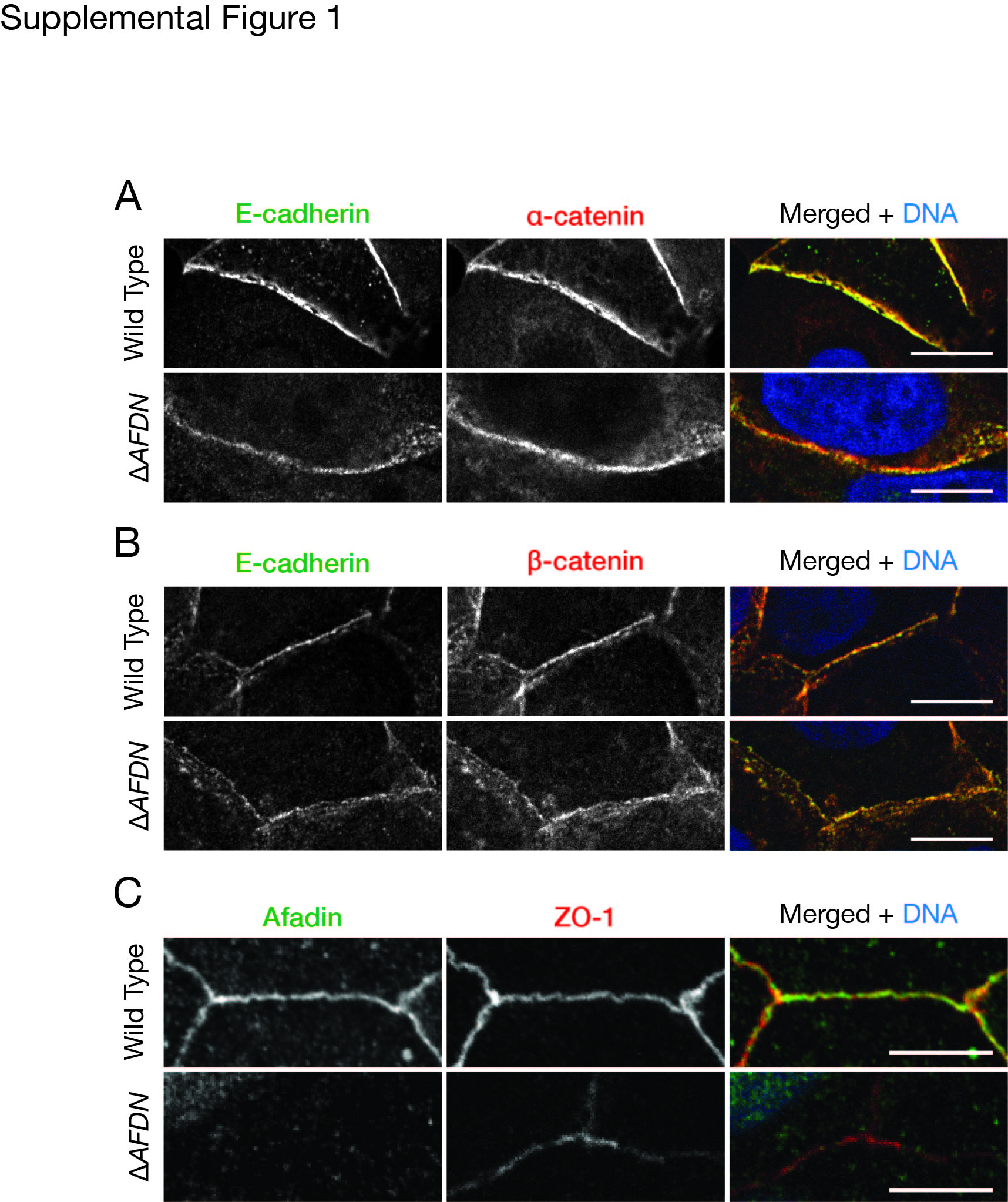

### Supplemental Figure 2

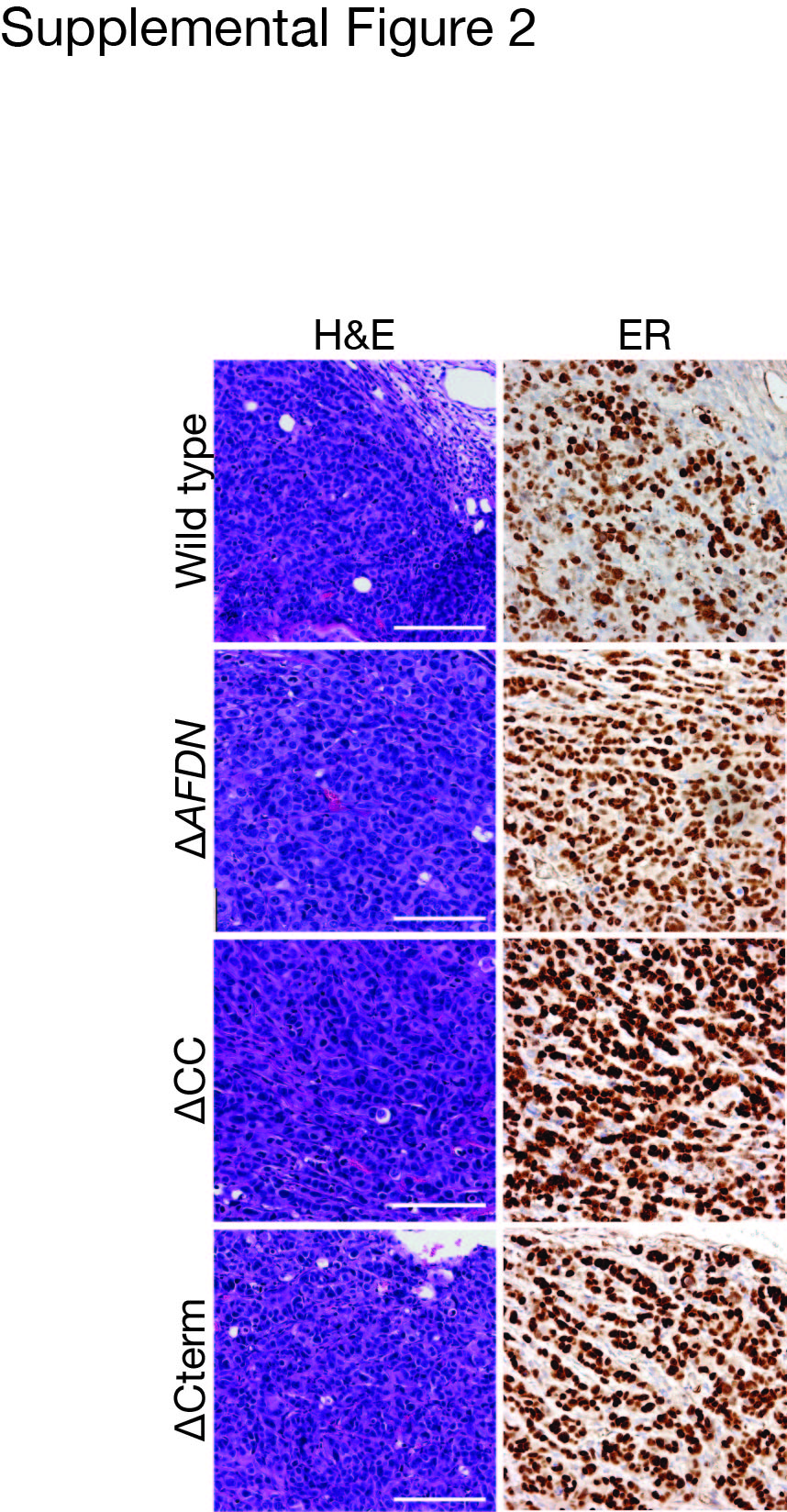

### Supplemental Table 1

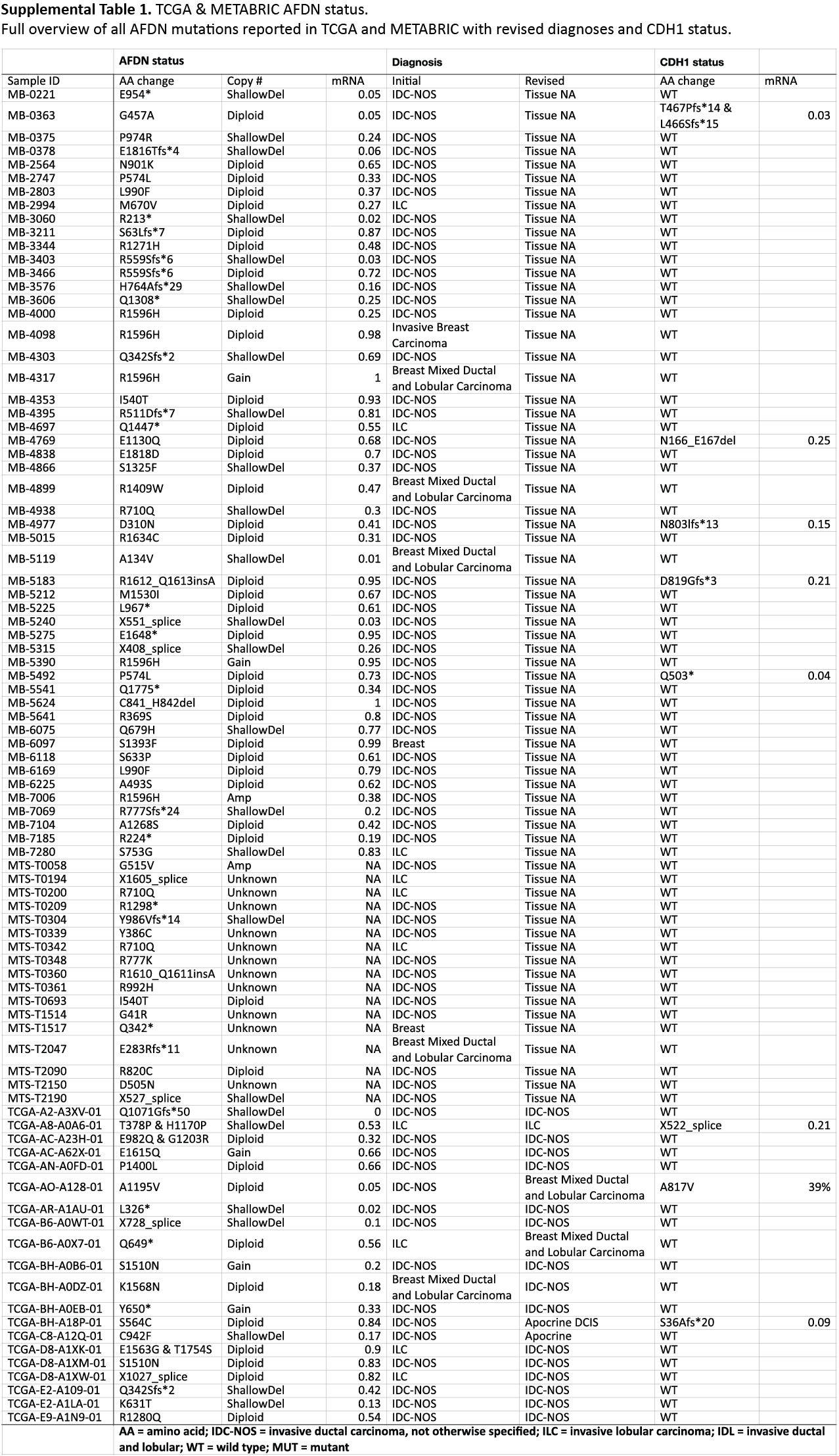

### Supplemental Table 2

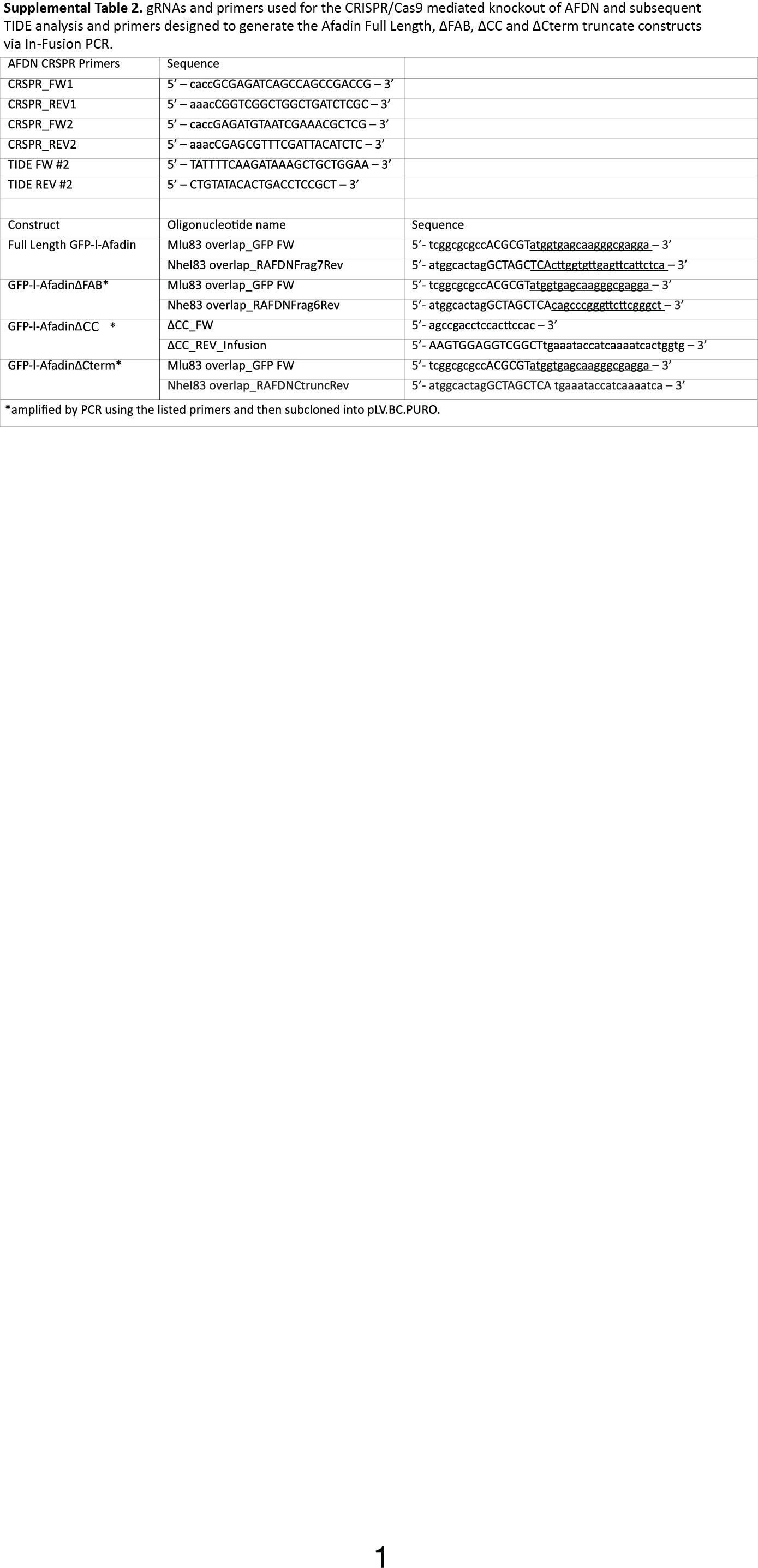
